# Nationwide multi-omics profiling of Japanese jack mackerel reveals geographic gut microbiome structuring despite host panmixia

**DOI:** 10.64898/2026.08.20.745924

**Authors:** Masa-aki Yoshida, Keito Tsunoda, Hirotoshi Kasane, Ayaka Kishimoto, Shunsuke Mori, Keito Komiya, Mayuko Hamada, Toshio Sekiguchi, Yoshiyuki Goto, Naoko Ishikawa, Yoshihisa Suyama, Davin H. E. Setiamarga

## Abstract

Host genetic markers often fail to resolve regional origins in highly connected or panmictic marine species. The Japanese jack mackerel, *Trachurus japonicus*, is a commercially important fishery species around Japan that shows little or no detectable population structure. Here, we used nationwide multi-omics profiling to compare host genomic variation and gut microbiome composition in wild *T. japonicus* collected from coastal regions across Japan. We generated MIG-seq data for 43 individuals and 16S rRNA gene profiles for 24 individuals; after quality filtering, 19 individuals remained for matched host–microbiome comparison. Genome-wide host SNP analyses showed weak or absent geographic population structure, consistent with previous evidence of panmixia in Japanese waters. In contrast, gut microbiome composition showed geographic structuring based on Bray-Curtis dissimilarity and PERMANOVA, and this pattern was not explained by proximity to river mouths or host-related variables. Locality- or individual-associated bacterial lineages contributed to the observed differences in the microbiome, while chloroplast-associated and Cyanobacteria-assigned ASVs suggested recent dietary or environmental input. These results indicate that gut microbiome can show regional biological variation not apparent from host genetic markers alone. Our study provides a proof-of-concept example of integrating host genomics and gut microbiome profiling to evaluate regional characteristics and origins in highly connected marine animals.

**Importance:** Highly dispersive marine fishes can remain genetically homogeneous across broad regions while encountering strongly contrasting environments. This disconnect limits the ability of host genetic markers to detect regional biological variation and raises a fundamental question: can host-associated microbial communities retain spatial ecological information when host population structure is weak? Using Japanese jack mackerel collected across Japan, we found weak or absent geographic structuring in host SNPs but significant locality-related variation in gut microbiome composition. The microbial pattern included broadly distributed marine-fish-associated taxa together with locally or individually enriched lineages and dietary or plankton-associated signals. These findings show that host-associated microbiomes can respond to regional ecological exposure at spatial scales not resolved by host population genomics. Beyond its potential for seafood traceability, this study establishes a framework for examining how environmental heterogeneity is recorded in animal-associated microbiomes under high host gene flow.

## Introduction

The coastal and offshore waters around the Japanese archipelago extend from cooler northern to warmer southern waters and are shaped by major current systems, including the warm Kuroshio and Tsushima Warm Currents and the cold Oyashio Current (Takikawa et al., 2005; Sakurai, 2007; Andres et al., 2008). These currents affect seawater temperature, productivity, larval transport, and coastal habitat conditions, creating environmental contrasts among the Pacific coast, the Sea of Japan, the Seto Inland Sea, and northern coastal waters (Itoh and Kimura, 2007; Kuroda et al., 2020; Wang et al., 2022). This hydrographic complexity is closely related to Japan’s highly diverse coastal ecosystem, marine biodiversity, and fisheries productivity, with more than 33,000 marine species recorded from Japan’s Exclusive Economic Zone, even when accounting for gaps in taxonomic and geographical knowledge (Fujikura et al., 2010; Yatsu et al., 2013; Nishikawa et al., 2020; Setiamarga et al., 2025a). The Marine Ecoregions of the World framework also places Japan within several temperate Northwest Pacific provinces, reflecting the biogeographic heterogeneity of the archipelago (Spalding et al., 2007). Recent large-scale environmental DNA surveys have also shown that coastal fish communities differ across Japan along environmental and biogeographic niche axes (Osada et al., 2026).

Population genetic studies of coastal and shelf fishes in this region, depending on life history, dispersal, spawning behavior, and anthropogenic influences, have shown that species span a continuum from highly connected populations to locally structured units. At the intraspecific level, the diverse environmental conditions around Japan do not necessarily mean a complete absence of population genetic structuring in marine fishes (Miya and Nishida, 1997); however, strong dispersal capabilities and the passive transport of larvae by ocean currents can facilitate and maintain gene flow across vast geographical ranges. Consequently, any resulting population structure may be weakened, restricted to localized scales, or otherwise remain highly challenging to detect (White et al., 2010; Selkoe and Toonen, 2011; Riginos et al., 2011). In high-gene-flow marine fishes, detectable population differentiation may depend strongly on marker choice, analytical resolution, or loci associated with selection (André et al., 2011; Liu et al., 2016). In Japanese coastal and shelf fishes, weak regional divergence or localized subdivision has been reported in species such as red sea bream (Blanco Gonzalez et al., 2015), marbled flounder (Sato et al., 2018), and Pacific cod (Sakuma et al., 2019). Marine fishes, therefore, provide an informative system for studying population connectivity and spatial biological variation, because their genetic population structures can differ from fisheries-management units and can be difficult to define when dispersal and gene flow are high (Nielsen and Kenchington, 2001; Hauser and Carvalho, 2008; Reiss et al., 2009).

The Japanese jack mackerel, *Trachurus japonicus*, is a pelagic fishery species distributed in East Asian seas and managed in Japan under Pacific Japan and Tsushima Warm Current stock units (Kanaji et al., 2009; Sassa et al., 2016; Muko et al., 2023). Eggs and larvae of *T. japonicus* are transported from spawning areas in the southern East China Sea toward nursery and coastal recruitment areas by the Kuroshio-Tsushima current system (Kim et al., 2007; Kasai et al., 2008; Sassa et al., 2008), while recruitment is also affected by juvenile growth, oceanographic variability, and transport processes (Takahashi et al., 2022), including interannual variation in the Kuroshio-Tsushima current system (Igeta et al., 2023). Earlier population genetic studies based on mitochondrial DNA (Song et al., 2013) and AFLP (Zhao et al., 2015) suggested that *T. japonicus* is a highly connected marine fish with little or no detectable population structure across Japanese to Chinese coastal waters. A recent genome-wide study using 19,904 SNPs from 614 individuals sampled around Japan likewise found no detectable geographic population structure, providing strong support for panmixia in *T. japonicus* (Hirao et al., 2024). The studies above thus support a highly connected, approximately panmictic population across Japanese waters.

Although Japan has long been supported by an abundance of marine resources, recent environmental changes have begun to cast a shadow over their stability. Regional origin information is also relevant to Japanese food movements such as “*chisan-chisho*”, or local production for local consumption, which emerged in response to concerns about food-system trust, rural decline, and food scandals in Japan (Kimura and Nishiyama, 2008). Seafood traceability is increasingly important for verifying product origin, supporting food safety, improving supply-chain transparency, and reducing illegal or unsustainable fishing (Helyar et al., 2014; Lewis and Boyle, 2017; André, 2018; Setiamarga et al., 2025b). In fisheries management, genetic data can help identify biological populations, stock mixing, and conservation units, but this approach depends on the presence of spatial genetic signals that can be used to distinguish groups or regions (Ogden, 2008; Reiss et al., 2009; Hemmer-Hansen et al., 2019; Cusa et al., 2022). The limited geographic resolution of host genetic markers in *T. japonicus* indicates the need for additional biological layers to evaluate regional characteristics and origins within this fishery species.

Fish gut microbiome provide a candidate biological layer for evaluating regional characteristics and origins within highly connected marine fishes. Gut bacterial communities are shaped by both host-associated and environmental factors, including salinity, trophic level, host phylogeny, habitat, and feeding habit (Sullam et al., 2012; Huang et al., 2020). In marine fishes, gut-associated microbes are linked to nutrition, physiology, immunity, and host-environment interactions, yet baseline data from wild marine fishes remain limited compared with those from aquaculture or model systems (Ghanbari et al., 2015; Egerton et al., 2018; Talwar et al., 2018). Host genetic homogeneity does not necessarily imply uniformity in gut microbiome composition, because feeding opportunities, habitat use, and exposure to environmental microbial pools can vary across space even when population connectivity is high (Jones et al., 2018). Recent comparative studies have shown that fish gut microbiome composition varies across host taxa and habitats, and that habitat can sometimes explain gut microbial variation more strongly than host taxonomy or trophic level (Huang et al., 2020; Kim et al., 2021). These findings suggest that gut microbiome can show locality-related variation potentially reflecting ecological exposure (Sullam et al., 2012; Kim et al., 2021). However, geographically broad, single-species studies that integrate host genomic variation with gut microbiome composition remain rare, especially for wild marine fishes with little or no detectable host population structure (Egerton et al., 2018; Jones et al., 2018; Lilli et al., 2024). In *T. japonicus*, the limited geographic resolution of host genetic markers makes regional characteristics and origins difficult to evaluate from host data alone, whereas gut microbiome may offer an additional biological layer in this highly connected fishery species.

In this study, we used nationwide, parallel profiling between host genetic variation and gut microbiome composition in wild *T. japonicus* collected from coastal regions across Japan. We tested whether gut microbial communities show geographic structuring in a fishery species whose host genetic markers show little or no regional differentiation. By linking host panmixia to gut microbial variation, we evaluated the gut microbiome as a microecological layer that may reflect regional biological characteristics in a commercially important marine fish. We, therefore, compared host genomic variation and gut microbiome composition to test whether the two biological layers show concordant geographic patterns in a highly connected marine fish.

## Materials and Methods

### 2.1 Species and sampling design

From July to October 2024, wild Japanese jack mackerel (*Trachurus japonicus*) were collected through citizen science-based fishing activities (“Nationwide Jack Mackerel Survey 2024”; Figure 1). The survey was conducted at a total of 13 survey locations by 11 participants. Sampling covered multiple coastal regions of the Japanese archipelago to capture broad geographic variation. The sampling sites and sample identifiers were as follows: Tottori offshore (IDs 1–2), Usuki River mouth, Oita (3–4), near the Nagoya University Marine Station, Mie (5–9), Obama Bay, Fukui (11–13), Akahama, Otsuchi Bay, Iwate (14–16, 40–42), Minato offshore, Hyogo (17–19), Maruyama fishing port, Hyogo (20–22), Etomo Port, Muroran, Hokkaido (23–28), Ushimado, Setouchi, Okayama (29–31), Okura-kaigan, Akashi, Hyogo (33–35), Hamamatsu, near the University of Tokyo Fisheries Laboratory (36–38), Nagoya Port, Aichi (39), and near the Oki Marine Station, Okinoshima, Shimane (44–46). Except for those from Tottori, samples were mainly obtained through flasher-rig fishing (“*sabiki*”). Samples from Tottori were purchased at a supermarket to include commercially available fish. For each individual, total length, body weight, sex, and maturity status were recorded when available. Additional metadata, including sampling date, locality, and processing batch information, were retained for downstream analyses. After capture, fish were frozen and transferred using frozen shipping services. The samples were preserved in frozen storage at −80°C until DNA extraction. Intestinal samples for microbiome analysis were collected by squeezing the intestines of the samples. Muscle tissues of all individuals used in this study were preserved for SNP analysis. After collecting tissue samples for DNA analysis, the obtained Japanese jack mackerel specimens were fixed in formalin and stored with specimen IDs (voucher IDs) according to the common rules established for the Ocean Shot research project (Table S1).

**Figure 1.**
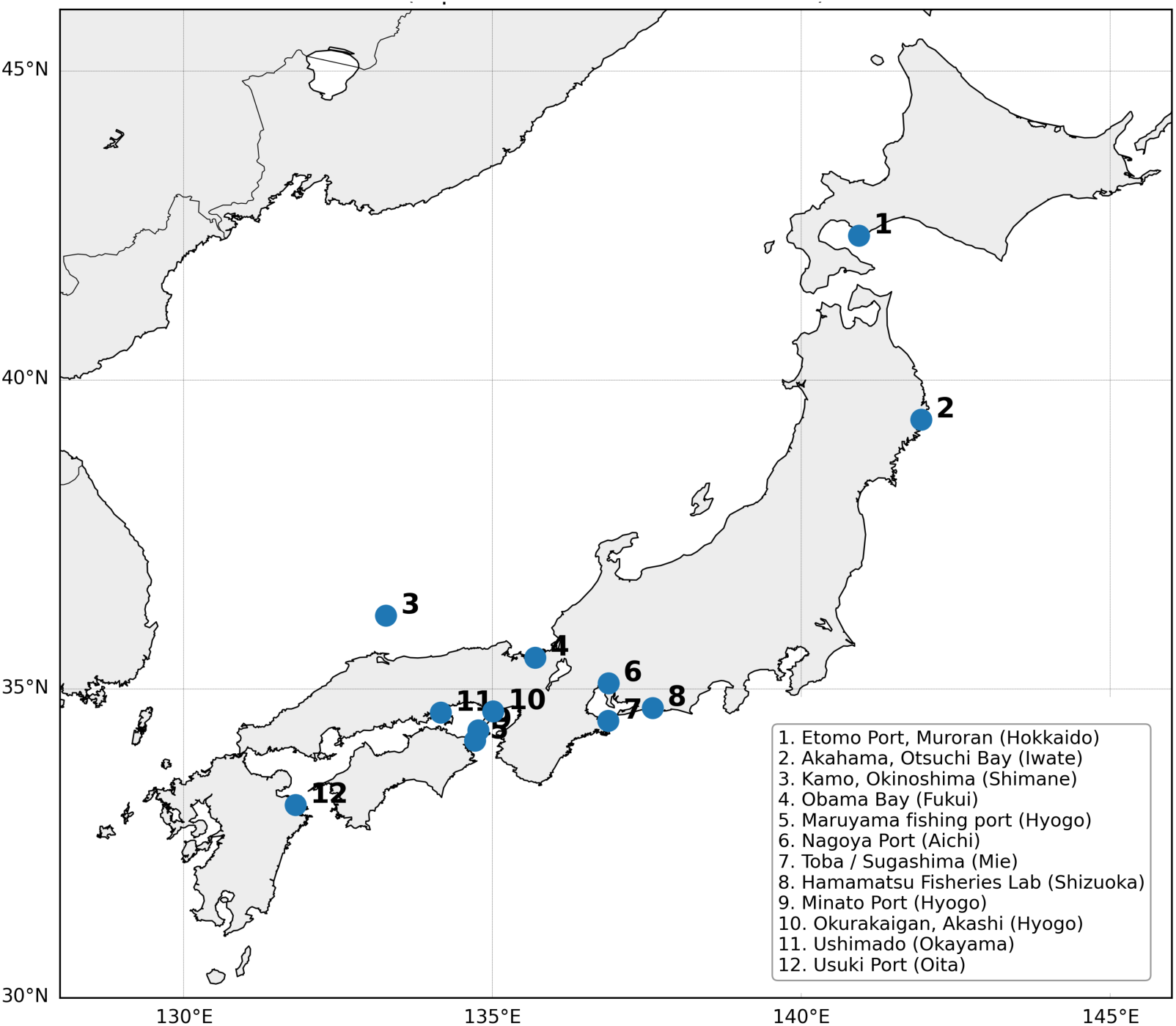
Sampling localities of Japanese jack mackerel (*Trachurus japonicus*) analyzed in this study. Numbered circles indicate representative coordinates of 12 named sampling localities across the Japanese archipelago. Site numbers correspond to the locality list shown in the figure. The base map was generated using the Basemap toolkit for Python, with coastline and geographic boundary data derived from the GSHHG/GMT datasets (Wessel and Smith, 1996).

### 2.2 Library construction and sequencing

#### 2.2.1 MIG-seq library construction and sequencing

Genome-wide SNP data were generated using multiplexed Inter-Simple Sequence Repeat (ISSR) genotyping by sequencing (MIG-seq), a PCR-based reduced-representation approach suitable for genome-wide SNP discovery and genotyping without restriction-enzyme digestion. The MIG-seq workflow consists of two PCR steps: a first multiplex PCR using tailed ISSR primers to amplify multiple inter-SSR regions from genomic DNA, followed by a second PCR to add Illumina adapter-binding sequences and sample-specific indices (Suyama et al., 2022).

Genomic DNA was extracted from the muscle using the QIAGEN Blood and Tissue Kit. DNA quantity and quality were assessed using the NanoDrop One. MIG-seq library preparation and sequencing were conducted following standard MIG-seq protocol (Suyama and Matsuki, 2015) but with minor adjustments. Briefly, the first PCR was performed with MIG-seq primer set-1 to amplify ISSR-flanking genomic regions. The second PCR was then conducted using indexed primers to append sequencing adapters and sample-specific barcodes. After amplification, individual libraries were pooled at approximately equimolar concentrations, purified, and size-selected to recover fragments in the range of approximately 300–800 bp. The pooled libraries were then quantified by qPCR and sequenced on an Illumina MiSeq platform using paired-end sequencing. To avoid the effects of low sequence diversity in the initial ISSR primer-derived bases, DarkCycle settings were applied during sequencing as described in the original MIG-seq protocol.

#### 2.2.2 16S rRNA gene library construction and sequencing

Gut microbiome composition was characterized by 16S rRNA gene amplicon sequencing. DNA was extracted from intestinal content using the NucleoSpin Microbial DNA Kit (Takara Bio), with bead beating (physical disruption) performed according to the protocol. DNA concentration and purity were evaluated using the NanoDrop One. We commissioned the Bioengineering Lab. Co., Ltd. to prepare an amplicon library targeting the V4 region of the 16S rRNA gene and perform sequencing. Sequencing was performed on an Illumina NextSeq 1000 platform using the NextSeq 1000/2000 P1 Reagents (600 Cycles) kit to generate 300 bp paired-end reads.

### 2.3 Data analysis

#### 2.3.1 SNP processing and population-genetic analysis

Raw MIG-seq reads were demultiplexed by sample index, and low-quality reads, primer-derived regions, and adapter-contaminated reads were removed using Trimmomatic v0.39 (Bolger et al., 2014) with the following parameters: ILLUMINACLIP (MIGadapter.fasta; 2:30:10), SLIDINGWINDOW:10:30, HEADCROP:6, CROP:77, and MINLEN:51. After trimming, forward (R1) and reverse (R2) reads were concatenated into a single FASTQ file for each sample.

The resulting high-quality reads were mapped to the *Trachurus japonicus* reference genome (Traja_1.0; GCA_045865225.1) using BWA-MEM v0.7.17 (Li, 2013). The resulting alignments were converted to BAM format and sorted using SAMtools v1.21 (Li et al., 2009). SNP discovery and genotyping were performed using gstacks implemented in Stacks v2.68 (Catchen et al., 2013; Rochette et al., 2019).

SNP datasets were generated using the populations module in Stacks. To evaluate the effect of locus occupancy on downstream analyses, loci were retained when present in at least 10% (R = 0.1) or 70% (R = 0.7) of sampled individuals. SNPs were filtered using a minimum minor allele count of three (–min-mac 3) and a maximum observed heterozygosity of 0.6 (–max-obs-het 0.6). Variant data were exported in PHYLIP format for phylogenetic analyses.

Maximum-likelihood phylogenetic analyses were conducted using IQ-TREE v2.3.6 (Minh et al., 2020). The best-fit substitution model was selected using ModelFinder (Kalyaanamoorthy et al., 2017), and ascertainment bias correction was implemented with the ASC option because the dataset contained only SNP sites. Branch support was assessed using 1,000 ultrafast bootstrap (UFBoot) replicates (Hoang et al., 2018) and 1,000 SH-like approximate likelihood ratio test (SH-aLRT) replicates (Guindon et al., 2010).

#### 2.3.2 Microbiome sequence processing

16S rRNA gene reads were processed in QIIME2 v2024.10 (Bolyen et al., 2019) to infer amplicon sequence variants (ASVs). After demultiplexing the reads using the fastx_barcode_splitter tool from FASTX-Toolkit (ver. 0.0.14), primer sequences were removed using fastx_trimmer from the same toolkit. Subsequently, we used sickle (ver. 1.33) to trim sequences with a quality score below 20, and discarded any sequences shorter than 130 bp along with their paired-end mates. The paired-end reads were merged using the FLASH (ver. 1.2.11; Magoč and Salzberg, 2011) script under the following conditions: 16S rRNA V4 (515F-806RB) expected fragment length 250 bp, read length 230 bp, and minimum overlap 10 bp. After removing chimeric and noisy sequences using the DADA2 plugin in QIIME2 (Callahan et al., 2016), representative sequences and an ASV table were generated. Taxonomic classification was performed by comparing the representative sequences with the 97% OTUs from the Greengenes database (ver. 13_8; McDonald et al., 2012) using the feature-classifier plugin. Samples with 5,000 or fewer reads after removing the host sequences (5/24) were not used in subsequent analyses. After host-sequence removal and quality filtering, the number of assigned reads per sample ranged from 5,763 to 38,709, with a median of 15,367 reads, indicating moderate variation in sequencing depth among retained microbiome samples. Community composition and diversity were evaluated without rarefaction, using normalized abundance-based approaches and diversity metrics less sensitive to uneven read depth.

#### 2.3.3 Community-level statistical analysis

Community-level analyses of the gut microbiome were conducted using the ASV abundance table after removal of host-derived and low-depth samples. ASV counts were converted to relative abundances for each sample, and Bray–Curtis dissimilarities were calculated to evaluate differences in community composition among individuals. Non-metric multidimensional scaling (nMDS) was used to visualize among-sample variation in gut microbiome structure.

To test whether gut microbial community composition differed among sampling localities or broader geographic categories, we performed permutational multivariate analysis of variance (PERMANOVA) using Bray–Curtis dissimilarity matrices with 999 permutations. The primary explanatory variables were sampling locality, broad marine region, and distance category from the nearest river mouth. Sampling date was also examined after transformation into sine and cosine terms to account for annual cyclicity. Where metadata were available, host-related variables such as total length, body weight, sex, and maturity status were also examined as candidate covariates. Because of the limited sample size and unbalanced sampling among localities, each explanatory variable was first tested individually, and multivariable models were interpreted cautiously.

All statistical analyses were performed using R (version 4.3.2) with the random seed set to 123 (set.seed(123)). For ordination (metaMDS and envfit functions), PERMANOVA (adonis2 function), and dbRDA (capscale, anova, and RsquareAdj functions), the ‘vegan’ package (version 2.6.4; Oksanen et al., 2022) was used. Hierarchical clustering was performed using the base hclust function, and Renyi diversity profiles were computed with the renyi function. Pairwise PERMANOVA was conducted using the pair-wise.adonis function from the ‘pairwiseAdonis’ package (v0.4.1; Arbizu 2020) with 999 permutations. Visualizations of nMDS ordinations and diversity profiles were generated using the ‘ggplot2’ package (v3.5.1; Wickham 2016).

## Results

### 3.1 Nationwide sampling design and overview

*Trachurus japonicus* were collected from 13 coastal localities across the Japanese archipelago between July and October 2024 (Figure 1; Table S1). Sampling covered a broad latitudinal range of their habitat, from Hokkaido to Kyushu. Most individuals were obtained by local angling, whereas the Tottori samples were purchased from a supermarket and are therefore treated with caution in interpretation.

A total of 43 individuals were included in the study and had usable metadata on sampling locality, sampling date, and all voucher specimens are available (Table S1). All individuals were retained for MIG-seq analysis. While 24 individuals were utilized for 16S rRNA-based gut microbiome analysis, sufficient reads were not obtained from 5 cases, and 19 individuals were used for the paired host–microbiome comparison (Table S1). This dataset thus represents a nationwide snapshot of genomic and gut microbiome variation in a single widespread marine fish species.

### 3.2 Genome-wide SNP data revealed weak or absent geographic population structure

Raw MIG-seq sequencing generated paired-end 80-bp reads for 43 individuals. Sequencing depth varied among individuals, ranging from 70,732 to 278,486 paired reads per sample, with a mean of 174,016 paired reads. The total sequence yield per individual ranged from 11.32 to 44.56 Mb. At the file level, Q20 and Q30 scores ranged from 95.43 to 98.35% and from 88.23 to 94.36%, respectively, and GC content ranged from 48.84 to 50.90%. These values indicate that the MIG-seq libraries provided broadly sufficient and comparable raw sequence quality for downstream SNP discovery and population-genetic analyses.

After reference-based SNP calling and filtering, 2,594 SNPs were retained using a stringent locus occupancy threshold (R = 0.7), whereas 10,444 SNPs were retained under a relaxed threshold (R = 0.1). Genome-wide SNP analyses revealed little evidence of geographic population structure among sampling localities. Maximum-likelihood phylogenetic analysis of the R = 0.7 dataset did not recover geographic clustering of individuals (Figure 2). To evaluate the robustness of this result, we repeated the analysis using the larger R = 0.1 dataset (Figure S1). Despite the fourfold increase in SNP number, individuals from the same locality still failed to form geographic clusters. Together, these results indicate that the MIG-seq dataset did not reveal clear geographic genetic structure across sampling regions.

**Figure 2.**
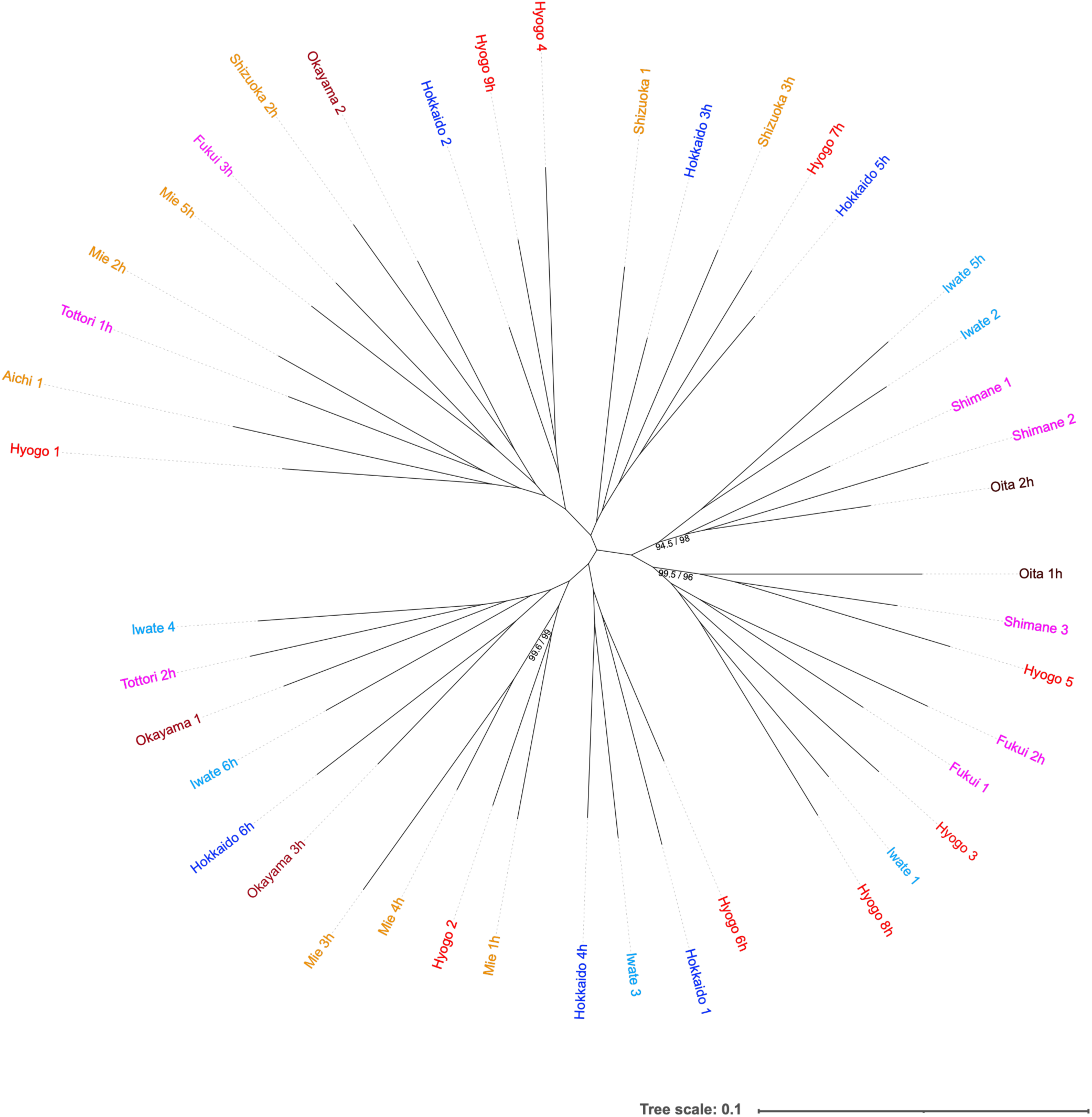
Maximum-likelihood phylogenetic tree of 43 Japanese jack mackerel individuals inferred from 2,594 genome-wide SNPs obtained from MIG-seq data (Stacks populations, R = 0.7). Samples are colored according to sampling locality. Branch support values indicate SH-like approximate likelihood ratio test (SH-aLRT) and ultrafast bootstrap (UFBoot) support, respectively. Only branches with SH-aLRT ≥ 80% and UFBoot ≥ 95% are shown. No well-supported clades corresponding to geographic localities were recovered.

### 3.3 Sequencing depth and quality of the gut microbiome dataset

Raw 16S rRNA amplicon sequencing generated paired-end reads for 24 individuals. The number of reads was relatively even among samples, ranging from 48,340 to 49,868 paired reads per individual, and read lengths ranged from 35 to 301 bp. Across R1 and R2 files, the mean read length ranged from 213.7 to 290.4 bp. The total sequence yield per individual, calculated from paired files, ranged from 21.38 to 28.40 Mb. These data provided a broadly balanced starting point for downstream quality filtering, denoising, and ASV-based microbiome analysis.

These reads were grouped into 1,463 ASVs with 411,623 assigned reads. Of these, one (ASV_004) showed a 100% match with *Trachurus japonicus* isolate T.jap_BH02_03 small subunit ribosomal RNA gene, partial sequence (MH331261.1) in a web BLAST search (NCBI BLASTn, as of May 4th, 2026) and was excluded from subsequent analyses. Of the remaining ASVs, 1,343 were identified as bacterial in origin (matching the Greengenes database at 97%), while the others were considered to be likely of eukaryotic origin from the diet, accounting for 13.5% of all reads. Taxonomic assignment identified 26 bacterial phyla, with an additional 120 ASVs classified as Bacteria but unassigned at the phylum level (Figure 3A). At the phylum level, Proteobacteria was the dominant group, accounting for 46.9% of total reads and detected in all 24 samples, followed by Firmicutes (18.7%, 22/24 samples), Cyanobacteria (9.4%, 20/24), Actinobacteria (4.9%, 20/24), and Planctomycetes (2.8%, 19/24) (Figure S2). At the family level, Vibrionaceae was the most abundant classified lineage, accounting for 24.2% of total reads and occurring in 21 of 24 samples (Figure 3B). At the genus level, *Photobacterium* was the dominant classified genus, representing 19.6% of total reads and detected in 21 samples, followed by *Vibrio* (3.2%, 16/24 samples). While most ASVs were not identified at the species level, the ASV assigned to *Photobacterium damselae* was an exception, accounting for 6.0% of the total reads.

**Figure 3.**
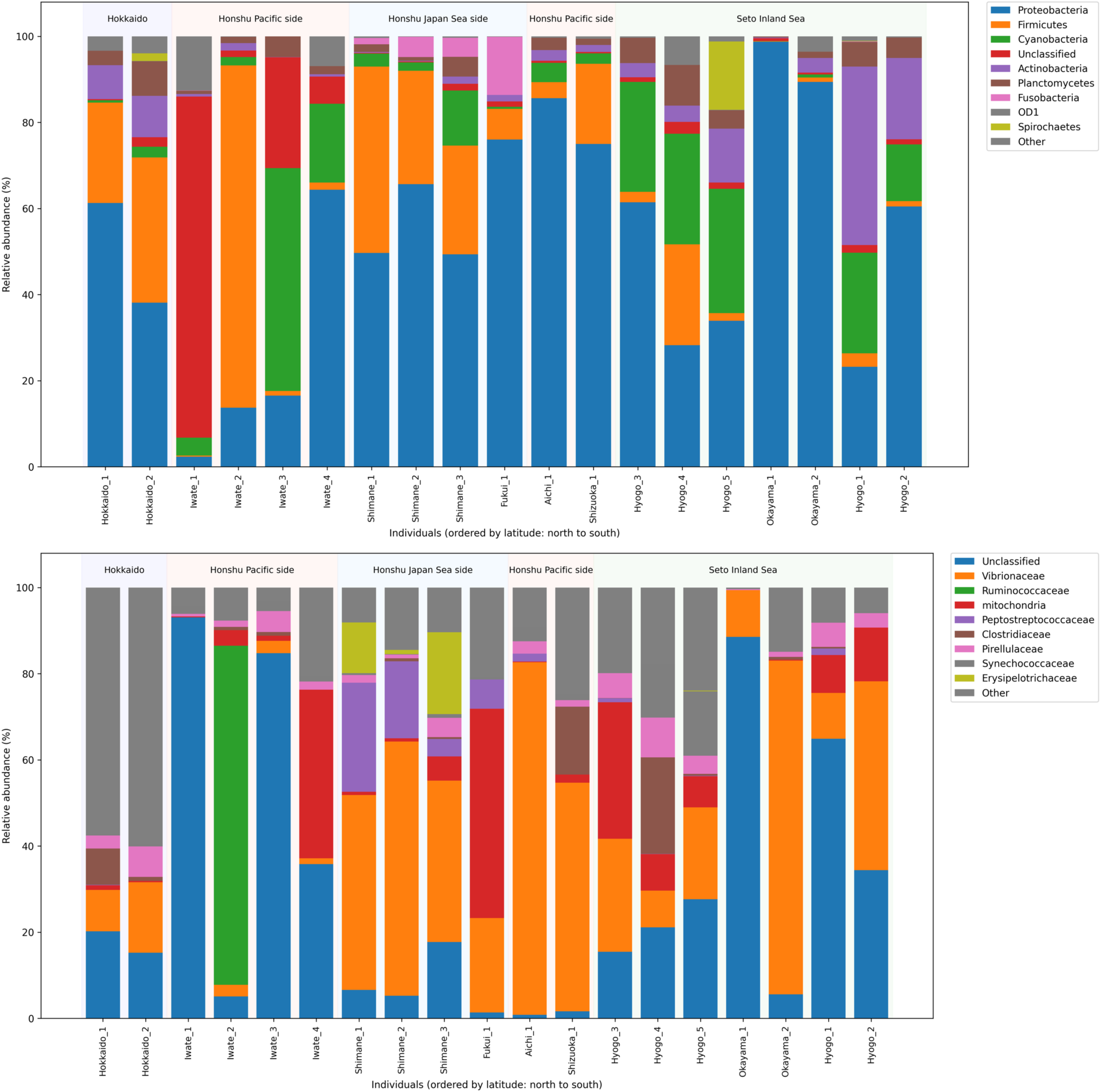
Relative abundance of the major bacterial phyla (A) and families (B) in the gut microbiome of individual *Trachurus japonicus*. Each bar represents one individual, ordered from left to right by sampling latitude (north to south). The nine most abundant phyla are shown explicitly; all remaining phyla are grouped as “Other” (gray). Background shading indicates the broad marine region of each sample: Hokkaido, Honshu Pacific side, Honshu Japan Sea side, or Seto Inland Sea.

Sequencing depth varied among individuals and observed ASV richness increased with read number (Figure S3), indicating that raw richness estimates were sensitive to library-size variation. Five samples with exceptionally low read counts and/or high proportions of host-derived reads were excluded from subsequent community analyses. These include samples from Tottori purchased at a supermarket. It should be noted that the results are affected by the amount and condition of the fecal matter. The remaining dataset provided sufficient coverage for comparative analysis of gut microbial community composition across localities.

### 3.4 Gut microbiome composition varied geographically across Japan

Hierarchical clustering suggested partial geographic grouping (Figure 4A). Non-metric multidimensional scaling (nMDS) based on Bray–Curtis dissimilarities revealed geographic structuring in gut microbiome composition (stress = 0.178; Figure 4). Individuals from the same locality tended to cluster together in ordination space, whereas samples from geographically distant sites were more dispersed (Figure 4B). PERMANOVA further indicated that locality explained a significant fraction of the variation in microbiome composition (R² = 0.24, p < 0.001), supporting the presence of broad-scale regional differentiation in the gut microbiome of *T. japonicus* (Table 1).

**Figure 4.**
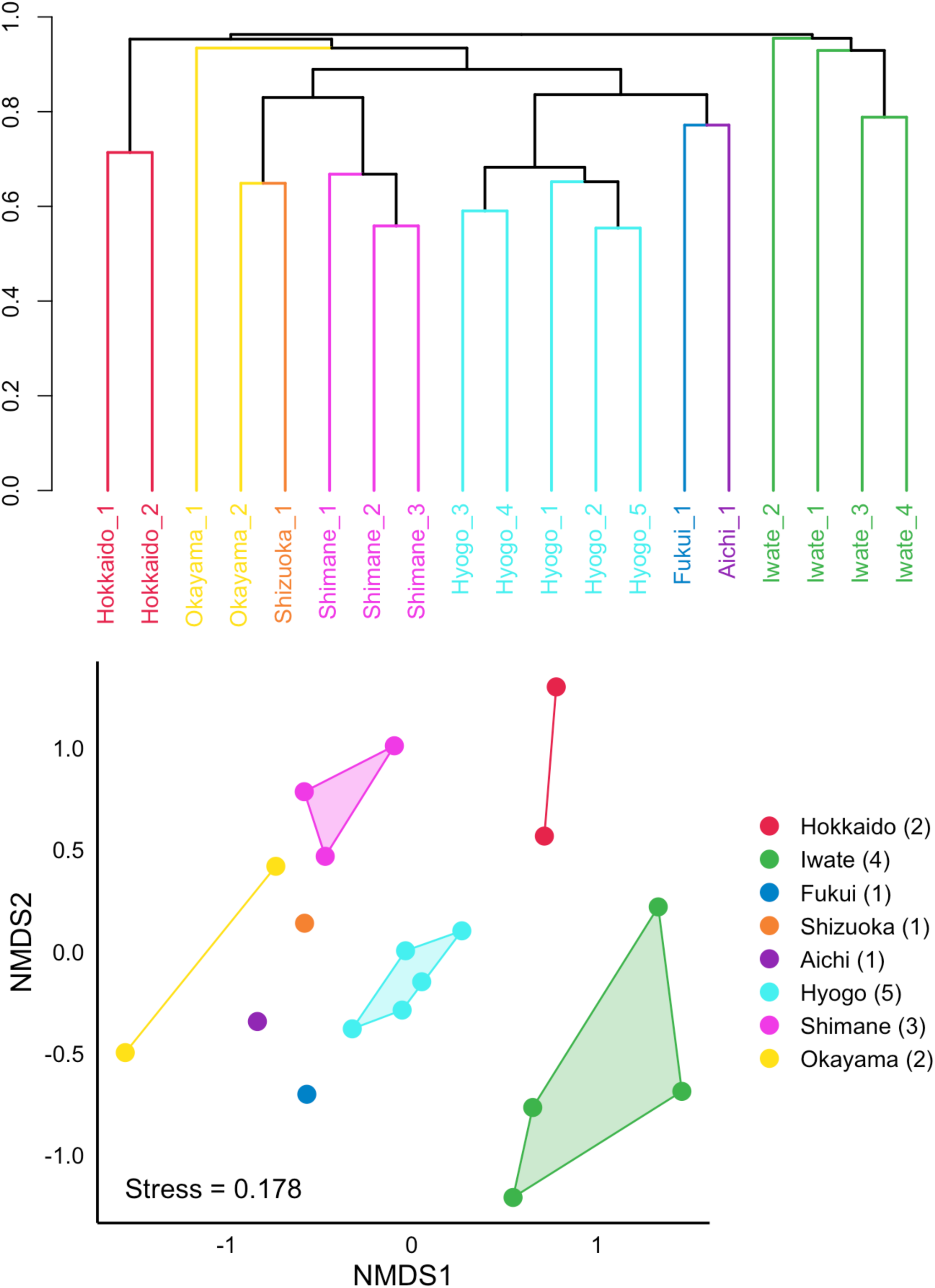
Geographic variation in the gut microbiome composition of Japanese jack mackerel. (A) Hierarchical clustering of 19 individuals based on Bray–Curtis dissimilarities calculated from ASV relative abundances. Sample-label colors indicate sampling locality. (B) Non-metric multidimensional scaling (nMDS) ordination based on the same Bray–Curtis dissimilarity matrix (stress = 0.178). Each point represents one individual, and colors indicate sampling locality. Individuals from the same locality tended to occupy similar positions, indicating a detectable locality-related pattern in gut microbiome composition.

**Table 1.**
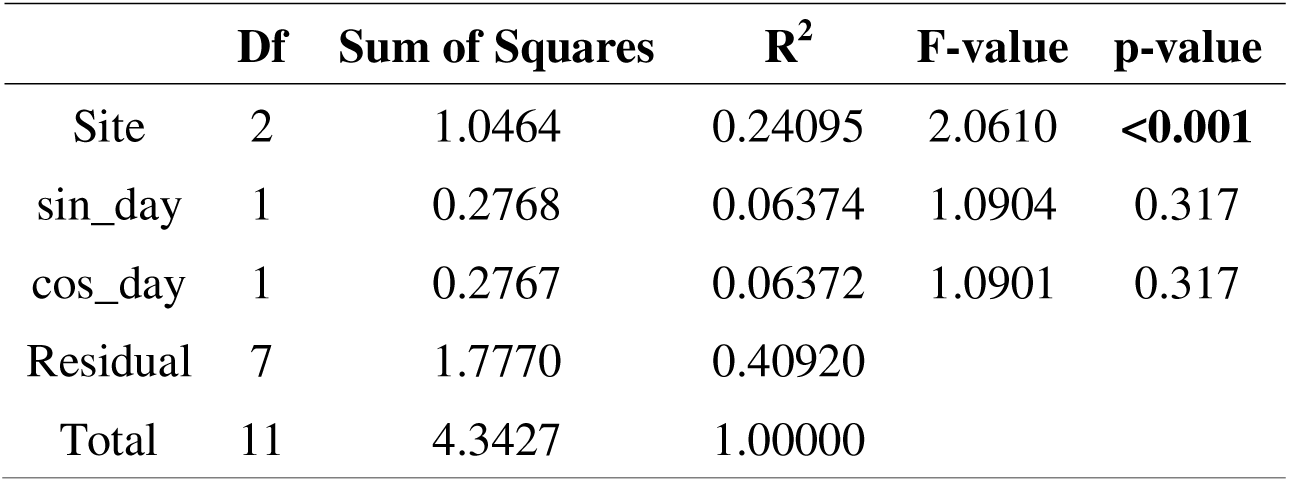
Permutational analysis of variance of gut microbial community composition by sampling site.

### 3.5 Region-specific bacterial lineages characterized local microbiomes

Although whole-community structure differed among localities, this pattern was also reflected in the distribution of specific bacterial lineages (Figure S4). Many of these locality enriched ASVs belonged to Proteobacteria and Firmicutes, including lineages within Vibrionaceae, Ruminococcaceae, Peptostreptococcaceae, Erysipelotrichaceae, and Fusobacteriaceae. Several taxa showed high total abundance but restricted prevalence across samples: Ruminococcaceae accounted for 6.5% of total reads but was detected in only 2 of 24 samples, whereas Erysipelotrichaceae and Fusobacteriaceae were detected in 4 and 7 samples, respectively. Peptostreptococcaceae accounted for 3.4% of total reads but occurred in fewer than half of the samples. These patterns indicate that part of the gut microbiome of *T. japonicus* consists of locally or individually enriched bacterial lineages, in contrast to broadly distributed taxa such as Vibrionaceae and Photobacterium. Because these taxa were not consistently detected across sampling sites, we treated them as locality- or individual-associated candidates rather than core components of the gut microbiome.

Beyond the influence of bacteria, it is also conceivable that the organisms consumed immediately prior to sampling played a role. For instance, in sample #14 (Iwate), unidentified reads (ASV_007) accounted for 71% of the total. Upon verification via web BLAST, these reads were estimated as an Alveolata-associated sequence. Such a finding may be considered characteristic of Japanese jack mackerel, given their planktivorous diet and relatively short digestive tracts. Furthermore, Cyanobacteria-assigned ASVs were also detected in several individuals, but most abundant ASVs within this phylum were annotated as chloroplast-associated sequences rather than clearly bacterial cyanobacteria. For example, ASV_012, classified as chloroplast-derived, accounted for 43.1% of assigned reads in sample #40 (Iwate), whereas several Chlorophyta- or Stramenopiles-associated chloroplast ASVs were detected in subsets of individuals. In contrast, ASVs assigned to *Synechococcaceae*/*Synechococcus* were more broadly detected, with ASV_018 occurring in 13 of 19 individuals and reaching 7.3% in one sample (#35, Hyogo). These patterns suggest that the Cyanobacteria-assigned fraction of the dataset likely reflects a mixture of marine picocyanobacteria and chloroplast-derived dietary or planktonic signals rather than a uniformly resident gut bacterial component.

## Discussion

### 4.1. Gut microbiome profiling as an approach for geographic traceability

Regional biological variation in highly connected marine fishes can be difficult to assess from host genetic markers alone, because high dispersal potential and gene flow often produce weak or absent population genetic structure in these fishes (Poulsen et al., 2011; André, 2018; Gagnaire et al., 2015). Broad connectivity and the resulting lack of geographic genetic structure limit the use of host genetic markers to resolve regional characteristics and origins (Manel et al., 2005; Ogden and Linacre, 2015; Powell and Campbell, 2020), thereby making DNA-based traceability difficult (Cusa et al., 2022).

The absence of spatial genetic differentiation, however, does not necessarily mean that individuals from different regions experience the same ecological conditions (Tian et al., 2014; Yatsu, 2019; Itsukushima, 2023), because ecological differences can persist even when population-genetic structure is weak or absent (Limborg et al., 2012; Diopere et al., 2018). These ecological differences among regions may affect phenotypes or characteristics that could be useful as identification markers for localities (Swain and Foote, 1999; Campana, 2005; Catalano et al., 2014). Regional genetic signals may also be blurred by anthropogenic gene flow associated with stock enhancement or aquaculture escapees, because hatchery- or farm-derived individuals can introgress into wild populations and modify genetic variation independently of natural dispersal (Blanco Gonzalez et al., 2015; Sawayama et al., 2026). The gut microbiome offers one way to examine this mismatch because microbial communities can respond to host-associated and environmental inputs over shorter timescales than host population-genetic differentiation (David et al., 2014; Petersen et al., 2023). Some studies have explored microbiome-based approaches for geographic-origin inference in marine animals, such as gut bacterial 16S rDNA fingerprinting of cultured seabass as a tracing tool (Pimentel et al., 2017), NGS-generated microbiome data and machine learning for Manila clam traceability (Milan et al., 2019), raw clam microbiomes as markers of harvest location (Liu et al., 2020), mussel gut microbiota fingerprints for geographic-origin tracing (del Rio-Lavín et al., 2023), gill microbiomes for distinguishing mackerels from the Atlantic and the Mediterranean (Piredda et al., 2023), and targeted metagenomics of gill bacterial communities for detecting site-related signatures in wild-caught seabass and seabream (Meriggi et al., 2024).

The Japanese jack mackerel, *Trachurus japonicus*, is an important pelagic fishery species managed in Japan under broad stock units (Kanaji et al., 2009; Sassa et al., 2016; Muko et al., 2023). Previous studies have repeatedly suggested high genetic connectivity across Japanese and adjacent East Asian waters (Song et al., 2013; Zhao et al., 2015; Hirao et al., 2024). In addition, its early-life transport through the Kuroshio-Tsushima current system provides a biological basis for broad connectivity (Kim et al., 2007; Kasai et al., 2008; Sassa et al., 2008; Igeta et al., 2023). In this study, we compared host genomic variation and gut microbiome composition in wild *T. japonicus* collected across coastal Japan to test whether these two biological layers show concordant geographic patterns. Our results indicate that gut microbiome composition retained a detectable locality-related signal despite the weak or absent geographic structuring of the host’s SNPs, thus suggesting that gut microbiome profiling can reveal locality-related biological variation that is not resolved by host genomic markers alone.

### 4.2. Geographic structuring of gut microbiota despite host panmixia

The central contrast emerging from our comparison is that host SNPs and gut microbiome composition did not show the same geographic resolution. Genome-wide SNP analyses showed weak or absent geographic population structure in *T. japonicus*, with individuals from different coastal regions largely overlapping in SNP-based ordination and no discrete regional groups recovered by population-structure analysis. This result is consistent with previous mitochondrial DNA, AFLP, and genome-wide SNP studies of *T. japonicus*, which also found little or no detectable geographic differentiation across Japanese and adjacent East Asian waters (Song et al., 2013; Zhao et al., 2015; Hirao et al., 2024). The agreement among these marker systems is important because it indicates that the weak geographic structure observed here is not specific to MIG-seq or to a single analytical workflow. It is instead consistent with the biology of a pelagic fish whose early life stages are transported through the Kuroshio-Tsushima current system from spawning areas toward nursery and recruitment areas, a process that can maintain broad connectivity among coastal regions even across environmentally heterogeneous waters (Kim et al., 2007; Kasai et al., 2008; Sassa et al., 2008; Igeta et al., 2023). Host genetic markers therefore provided little geographic resolution in *T. japonicus* at the spatial scale examined here, establishing the background against which the gut microbiome result should be interpreted.

Meanwhile, gut microbiome composition showed geographic structuring across coastal Japan. Bray-Curtis nMDS indicated that individuals from the same locality tended to cluster together, while samples from geographically distant sites were more dispersed, and PERMANOVA indicated that locality explained a significant fraction of microbiome variation (R² = 0.24, p < 0.001). This pattern is notable because it shows that host genomic homogeneity did not correspond to uniformity in host-associated microbial communities. Comparable geography-associated gut microbiome patterns have been reported in other fishes, but their interpretation depends strongly on the host species, habitat, life history, and sampling design. For example, geographically isolated wild populations of the range-shifting marine rabbitfish *Siganus fuscescens* showed distinct hindgut microbial communities while retaining a shared core microbiome (Jones et al., 2018). In Mediterranean scorpionfishes *Scorpaena* spp., geographic origin and host phylogeny were both associated with gut mucosal microbiota, with geographic origin influencing microbial diversity and composition (Lilli et al., 2023). In two-banded sea bream, the gut microbiome was examined to test whether microbial community structure reflected geographic origin at large and small spatial scales (Lilli et al., 2024). These studies show that geographic structuring of fish gut microbiome is not unique to *T. japonicus*, thus supporting our present result. However, our present study provides an important step forward because the comparison was conducted within a single wild pelagic species that showed weak or absent host SNP structure.

The river-mouth analysis further refined the interpretation of the microbiome pattern. Distance category from the nearest river mouth did not explain gut microbiome composition in *T. japonicus* (R² = 0.04, p = 0.537, Figure S5), indicating that the locality-related signal was not simply a freshwater-influence effect. This result instead points to broader regional ecological differences, such as differences in water masses, prey fields, habitat use, and local microbial pools. Studies of wild and commercial fishes support the interpretation that locality-related differences in gut microbiome can reflect variation in recent feeding, surrounding water, habitat use, or local microbial pools. In adult wild fishes from the Yangtze River, water-derived microbes were inferred to enter the gut, and feeding habits contributed to similarities in gut microbial communities among fishes (Yang et al., 2022). In shallow-lake fishes, vertical habitat preference was the main factor associated with gut microbiome, exceeding the effects of host taxonomy and trophic level (Zhang et al., 2024). In commercial marine fishes from Fujian Province, gut microbiome also varied across host and geographic factors, with a persistent core microbiota detected across multiple scales (Sun et al., 2022). Although these studies do not provide direct analogues for *T. japonicus*, they support the interpretation that the locality-related microbiome signal observed here reflects differences in recent feeding, surrounding water, habitat use, or local microbial pools among coastal regions.

The contrast between the host SNP and gut microbiome results therefore reflects a difference in the biological information captured by the two layers. Host genomic variation in *T. japonicus* appears to be strongly influenced by dispersal and gene flow, which blur geographic differentiation across Japanese waters. The gut microbiome composition, by contrast, retained a locality-related signal that was not explained by simple river-mouth proximity and may reflect regional ecological conditions. This does not mean that the microbiome pattern can already be treated as a validated tool for locality identification. Rather, it shows that gut microbiome profiling can detect regional biological variation that host SNP markers did not resolve in this highly connected fishery species.

### 4.3. Shared and locality-associated components of T. japonicus gut microbiome

The gut microbiome of *T. japonicus* comprised a combination of broadly distributed marine-fish-associated taxa and more unevenly distributed locality- or individual-associated lineages. Our results showed that Proteobacteria, detected in all 24 samples, was the most abundant phylum, followed by Firmicutes, Cyanobacteria, Actinobacteria, and Planctomycetes. Meanwhile, Vibrionaceae was the most abundant family, and *Photobacterium* was the dominant genus, with both detected in most individuals. This composition is consistent with marine fish gut microbiome studies showing that salinity, trophic ecology, host phylogeny, and habitat are important correlates of fish gut bacterial communities, and that marine fishes often harbor Proteobacteria- and Vibrionales-rich gut communities (Sullam et al., 2012; Egerton et al., 2018; Kim et al., 2021). The high abundance and broad occurrence of *Photobacterium* in *T. japonicus* therefore should not be interpreted as a pattern unique to Japanese jack mackerel. In migrating Atlantic cod, shotgun metagenomic analysis showed that Vibrionaceae dominated fecal metagenomic reads, with *Photobacterium* representing most of this family (Le Doujet et al., 2019). This comparison supports the interpretation that the dominance and broad occurrence of Proteobacteria, Vibrionaceae, and *Photobacterium* observed here represent a recurrent marine-fish-associated gut microbiome component rather than a species-specific anomaly.

However, these broadly distributed taxa represented only one component of the gut microbiome. Several locality- or individual-associated ASVs belonged to Proteobacteria and Firmicutes, including lineages within Vibrionaceae, Ruminococcaceae, Peptostreptococcaceae, Erysipelotrichaceae, and Fusobacteriaceae. Some of these lineages showed high total abundance but restricted prevalence across individuals. Ruminococcaceae accounted for 6.5% of total reads but was detected in only 2 of 24 samples, whereas Erysipelotrichaceae and Fusobacteriaceae were detected in 4 and 7 samples, respectively. Peptostreptococcaceae accounted for 3.4% of total reads but occurred in fewer than half of the samples. These patterns indicate that the gut microbiome of *T. japonicus* comprises a broadly distributed bacterial component alongside a more variable fraction enriched in particular individuals or localities. Since these taxa were not consistently detected across sampling sites, the contrast between broadly distributed taxa and locality- or individual-associated lineages is important for interpreting variations in among-sample microbiome (Neu et al., 2021; Lilli et al., 2024). The locality-related microbiome signal detected in this study probably does not arise from the dominant shared component alone, but from the uneven distribution of locality- or individual-associated lineages across samples.

The locality- or individual-associated fraction may reflect recent feeding history, prey-associated microbes, exposure to local seawater microbial pools, or other short-term ecological inputs. Diet and recent ingestion can shape fish gut microbiomes through dietary microbial input, prey-associated microbiomes, and transient food-associated communities (Smith et al., 2015; Escalas et al., 2021; Restivo et al., 2021; Viver et al., 2023). Geographic location can also be associated with gut microbiome in wild marine fishes, although these communities may remain distinct from surrounding water microbiota (Soh et al., 2024). The same short-term ecological signal was also apparent in the non-bacterial and plastid-associated sequences detected in the dataset. In sample #14 from Iwate, unidentified reads assigned by BLAST to Alveolata accounted for a large proportion of the total reads, suggesting that recently ingested planktonic organisms or prey-associated gut contents can strongly influence the gut profile of some individuals.

Cyanobacteria-assigned ASVs were also detected in several samples, but many abundant ASVs within this category were annotated as chloroplast-associated sequences rather than clearly bacterial cyanobacteria. For example, a chloroplast-derived ASV accounted for a large fraction of assigned reads in sample #40 from Iwate, and Chlorophyta- or Stramenopiles-associated chloroplast ASVs were detected in subsets of individuals. Because chloroplast sequences can originate from ingested algae, phytoplankton, or prey-associated gut contents, their uneven distribution among individuals probably records recent feeding or environmental exposure rather than stable colonization of the gut (Watanabe et al., 2021). The broader occurrence of Synechococcaceae/*Synechococcus* ASVs provides a useful contrast, because this fraction may include marine picocyanobacteria in addition to plastid-derived signals. The enrichment of chloroplast-associated ASVs in particular individuals supports the view that part of the gut microbiome profile reflects recent dietary or environmental exposure. Because chloroplast sequences can originate from ingested algae, phytoplankton, or prey-associated gut contents, their uneven distribution among individuals may record short-term variation in feeding history rather than stable colonization of the gut. While certain Cyanobacteria are known to be non-photosynthetic, it would be reasonable to infer that they entered the digestive system either directly from the surrounding waters or via prey such as copepods ingested shortly beforehand. In this sense, Cyanobacteria-assigned ASVs provide a useful contrast to broadly distributed marine-fish-associated taxa such as Photobacterium: the former may represent transient ecological input, whereas the latter may reflect a more persistent host- or habitat-filtered microbiome component.

### 4.4. Applied value and limitations of gut microbiome profiling for geographic traceability

Microbiome-based geographic-origin studies in marine animals have shown that microbial profiles can contain spatial information useful for traceability (Milan et al., 2019; Liu et al., 2020; del Rio-Lavín et al., 2023; Piredda et al., 2023). In this study, we directly compared host genomic variation and gut microbiome composition in a highly connected pelagic fish with weak host genomic structuring, consistent with previous evidence of panmixia. Despite the weak or absent geographic structure of the host species, gut microbiome composition retained locality-related variation. This contrast suggests that gut microbiome profiling can add a biological layer that host genomic markers alone may not capture, particularly in marine fishes where dispersal and gene flow reduce the power of genetic markers for regional-origin inference (Gagnaire et al., 2015; Zhang et al., 2020). Our results therefore provide a proof-of-concept for integrating host genomics and gut microbiome profiling in regional-origin research on panmictic pelagic fishery species such as *T. japonicus*. Further validation using independent test samples, explicit assignment models, and repeated sampling across seasons, years, and post-capture handling conditions will be needed before this approach can be developed into a practical traceability method.

Several limitations prevent the present microbiome signal from being treated as a validated traceability marker. First, the study represents a nationwide snapshot from a single sampling season and year. The samples were collected between July and October 2024, and the design does not separate spatial effects from seasonal or interannual variation. Second, the number of microbiome samples compared with the host SNP dataset was limited, with 24 individuals used for gut microbiome profiling and 19 individuals available for paired host-microbiome comparison. Third, sampling was uneven among localities, with one locality (from Tottori) included supermarket-purchased samples. Fourth, the microbiome signal was evaluated as a community-level locality effect rather than as a predictive origin-assignment model. For traceability use, future studies would need independent training and validation datasets, repeated sampling across seasons and years, and tests of whether microbiome profiles remain informative after differences in handling, storage, transport, and market distribution. This distinction is important because microbiome-based traceability studies that aim for applied origin assignment usually require explicit validation of classification performance and temporal stability, not only detection of geographic structure (Ogden, 2008; Lokmer et al., 2016; Duarte et al., 2025).

Our present 16S rRNA amplicon approach for microbiome analyses also limits the taxonomic and functional interpretation of the locality-related microbiome signal. Many ASVs could not be resolved to the species level, and 16S-based profiling cannot directly identify strain-level variation, functional genes, or metabolic activity (Poretsky et al., 2014; Heidrich and Beule, 2022). This matters because short amplicon data alone cannot confidently determine whether locality-associated ASVs reflect strain-level variation, transient dietary signals, environmental inputs, or functionally distinct bacterial lineages. Short-read 16S rRNA sequencing is useful for comparing community composition, but it has limited resolution for species- or strain-level identification compared with longer marker sequences or genome-level approaches (Johnson et al., 2019). Functional inference from 16S data is also indirect and should be interpreted cautiously, especially when ecological interpretation depends on the metabolic roles of particular bacterial lineages (Matchado et al., 2024). Future analyses could include shotgun metagenomics to help evaluate strain-level and functional differences among localities (Scholz et al., 2016; Rausch et al., 2019), and metatranscriptomics to test whether geographically associated microbial lineages also differ in active gene expression (Aguiar-Pulido et al., 2016; Bashiardes et al., 2016). Species- or strain-level identification of locality-associated taxa would also support the development of simpler and faster diagnostic tools, such as PCR-based barcoding (*e.g*., Walter et al., 2000; Wanka et al., 2018; Floris et al., 2021) and MALDI-TOF MS-based (*e.g*., Assis et al., 2017; Jlidi et al., 2022; Medina et al., 2024) microbial identification systems. These approaches would be especially useful for distinguishing resident gut bacteria from transient locality-related dietary or environmental sequences.

## Data availability

Raw sequencing reads were deposited in the Sequence Read Archive (BioProject IDs; MIG-seq DNA, PRJDB42559; microbiome DNA, PRJDB42560).

## Author Contributions (CRediT)

MAY, MH, TS, YG and DHES designed and conceived this study. HK and AK collected data. KT, SM, and KK analyzed and interpreted the results and drafted the manuscript. NI and YS supported statistical analyses. All authors read and approved the final manuscript.

## Acknowledgments

This study summarizes the outcomes of the Rinkai Hackathon 2025, a workshop held in collaboration with a research group called “RinkaiHack (https://sites.google.com/view/rinkaihack/).” We would like to express our gratitude to everyone who assisted in the planning and management of Rinkai Hackathon 2025. This study was also conducted with the support of JAMBIO (Japanese Association for Marine Biology).

The authors wish to extend their sincere gratitude to the following participants of the “2024 National Japanese Jack Mackerel Survey” for their invaluable contributions: Mr. Kiyoshi Saito (Oita), Mr. Kanta K. Ochiai (Mie), Mr. Hiroumi Ueno (Fukui), Dr. Jun Hayakawa (Iwate), Dr. Naotaka Hara (Hyogo), Ms. Kayo Takayama (Hokkaido), Mr. Takehiro Nishiyama, Ms. Kanoko Nishiyama (Kato) (Okayama and Hyogo), and Prof. Kiyoshi Kikuchi (Shizuoka). We also thank the students enrolled in the Department of Biotechnology at Nagoya Future Technical College during the 2024 academic year and their instructor, Dr. Takeshi Kawashima, for their assistance with the collection of the jack mackerel in Nagoya, Aichi Prefecture.

