## Supplementary material for "Nationwide multi-omics profiling of Japanese jack mackerel reveals geographic gut microbiome structuring despite host panmixia": Figure S

Supplemental figures

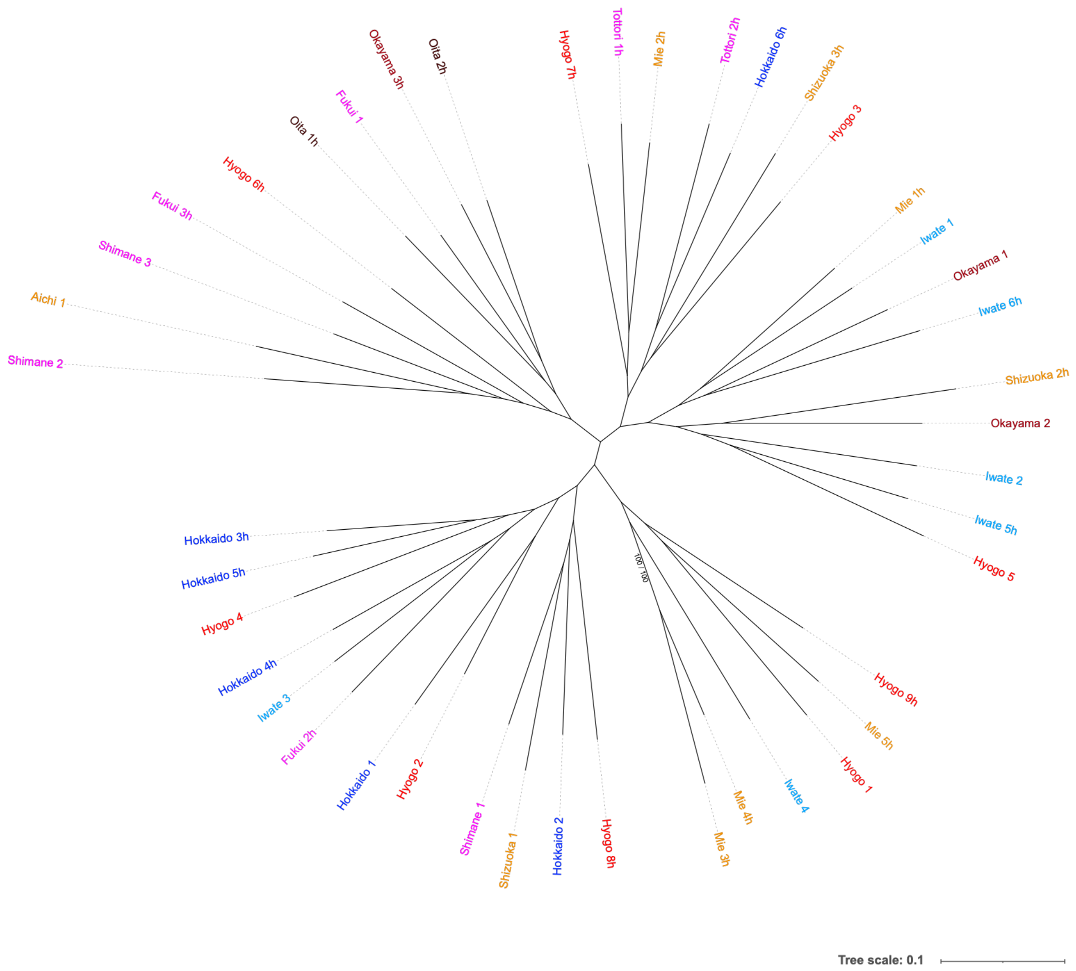


Figure S1. Maximum-likelihood phylogenetic tree of 43 Japanese jack mackerel individuals inferred from 10,444 genome-wide SNPs obtained using a relaxed locus-occupancy threshold (Stacks populations, R = 0.1). Samples are colored according to sampling locality. Branch support values indicate SH-aLRT and UFBoot support, respectively. Only branches with SH-aLRT ≥ 80% and UFBoot ≥ 95% are shown. Despite the substantially larger SNP dataset, no well-supported geographic clustering was detected.

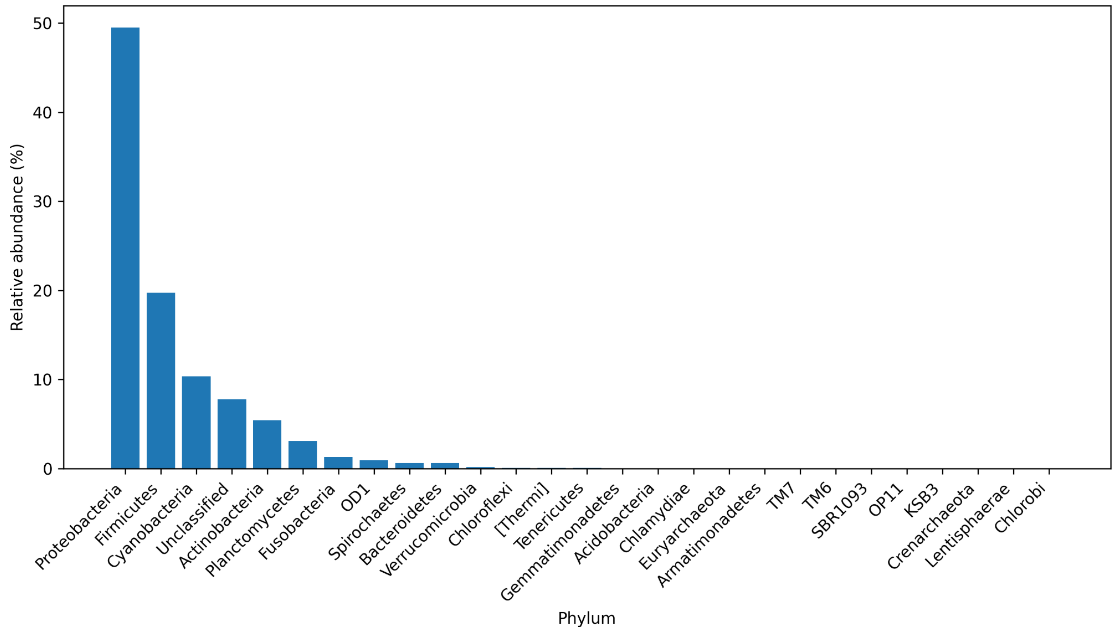


Figure S2. Overall phylum-level composition of the gut microbiome of Japanese jack mackerel (*Trachurus japonicus*). Relative abundance was calculated from pooled ASV read counts across the 19 individuals retained after host-sequence removal and read-depth filtering. Bars represent bacterial phyla in descending order of pooled relative abundance. ASVs classified as Bacteria but not assigned at the phylum level are shown as “Unclassified.”

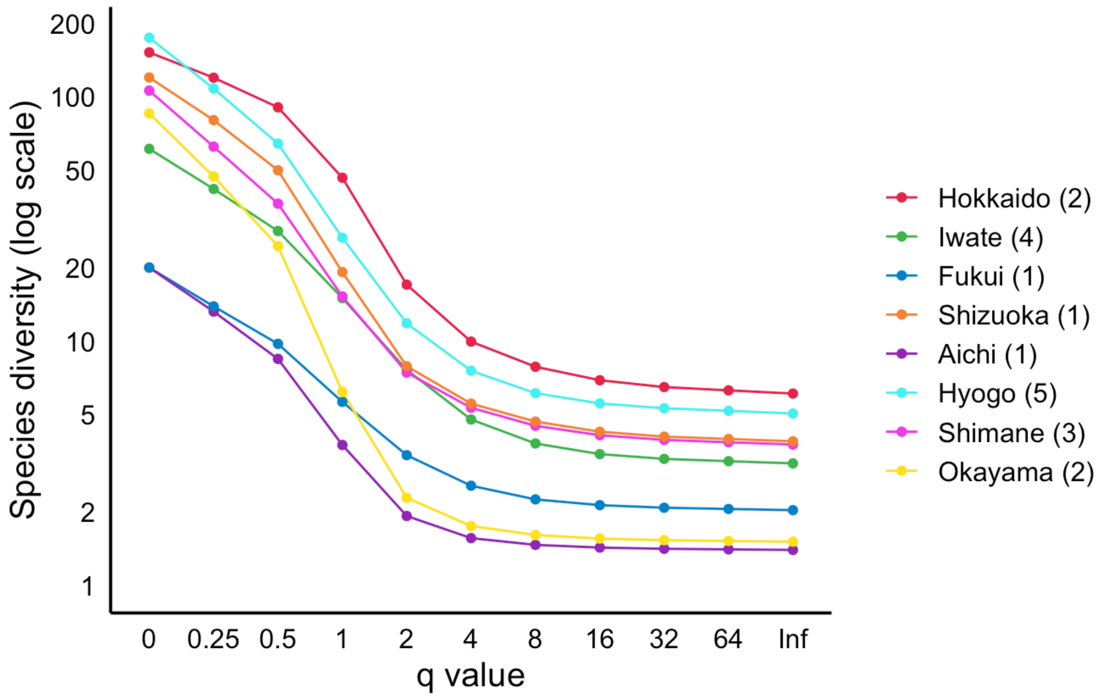


Figure S3. Rényi diversity profiles of gut microbiome samples from Japanese jack mackerel. Diversity is shown as a function of order q, with q = 0 corresponding to observed richness, q = 1 to the exponential of Shannon diversity, and q = 2 to the inverse Simpson index. Each line represents one retained individual and is colored according to sampling locality.

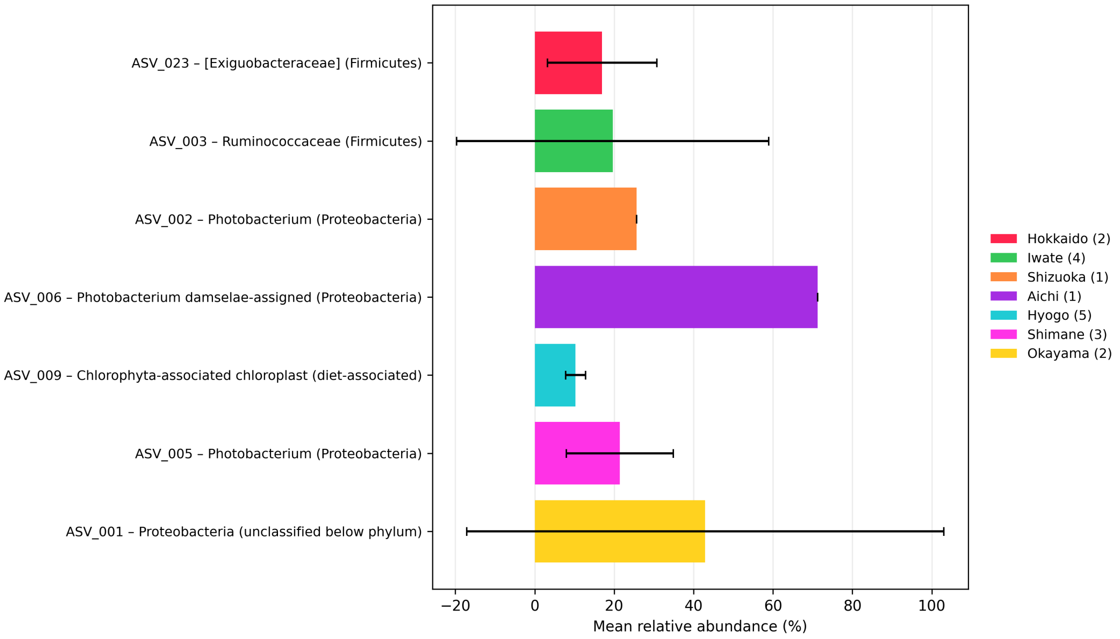


Figure S4. Locality- or individual-enriched bacterial and diet-associated ASVs in the gut contents of Japanese jack mackerel (*Trachurus japonicus*). Bars represent the mean relative abundance of selected ASVs within each sampling locality, calculated for each individual as the proportion of reads assigned to the ASV relative to the total assigned reads in that sample. Error bars indicate standard deviations among individuals within each locality; no error bar is shown for localities represented by a single individual. Colors indicate sampling localities, with the number of retained microbiome samples shown in parentheses. ASV identifiers and taxonomic assignments are shown at the lowest confidently assigned rank, with the corresponding phylum indicated in parentheses where applicable. ASV_009 was assigned to a Chlorophyta-associated chloroplast sequence and was retained as a putative diet-associated signal. Mitochondrial sequences were excluded from the figure.

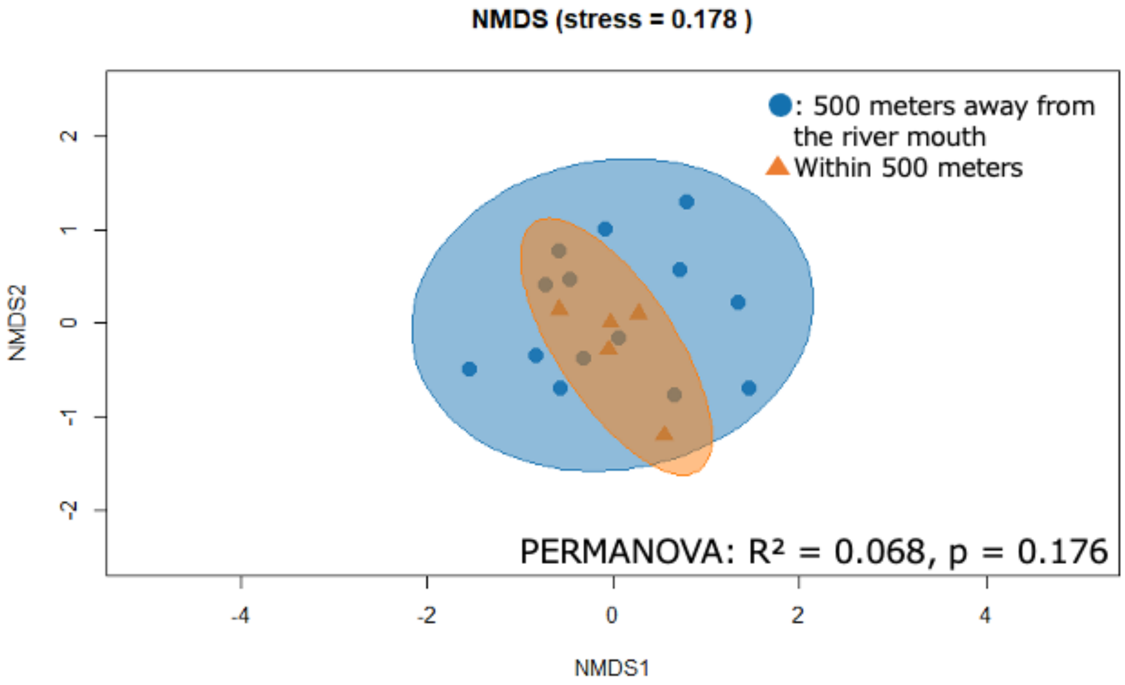


Figure S5. Gut microbiome composition in relation to proximity to river mouths. Non-metric multidimensional scaling based on Bray–Curtis dissimilarities are shown for the 19 retained individuals, classified according to whether the sampling locality was within or beyond 5 km of the nearest river mouth. Colors or symbols indicate river-mouth distance category. No clear separation was detected between categories, and PERMANOVA indicated no significant association between river-mouth proximity and community composition ($R^2 = 0.04$, $p = 0.537$).
